# An *α*-cyanostilbene derivative for the enhanced detection and imaging of amyloid fibril aggregates

**DOI:** 10.1101/2020.06.25.172627

**Authors:** Nicholas. R Marzano, Kelly M Wray, Caitlin L Johnston, Bishnu P Paudel, Yuning Hong, Antoine van Oijen, Heath Ecroyd

**Affiliations:** Molecular Horizons and School of Chemistry and Molecular Bioscience, University of Wollongong, Wollongong, NSW 2522, Australia; Illawarra Health & Medical Research Institute, Wollongong, NSW 2522, Australia; Department of Chemistry and Physics, La Trobe Institute for Molecular Science, La Trobe University, Melbourne, VIC 3086 Australia

## Abstract

The aggregation of proteins into amyloid fibrils has been implicated in the pathogenesis of a variety of neurodegenerative diseases, including Alzheimer’s and Parkinson’s disease. Benzothiazole dyes such as Thioflavin T (ThT) are well characterised and widely used fluorescent probes for monitoring amyloid fibril formation. However, existing dyes lack sensitivity and specificity to oligomeric intermediates formed during fibril formation. In this work we describe the use of an α-cyanostilbene derivative with aggregation-induced emission properties (called ASCP) as a fluorescent probe for the detection of amyloid fibrils. Similar to ThT, ASCP is fluorogenic in the presence of amyloid fibrils and upon binding and excitation at 460 nm produces a red-shifted emission with a large Stokes shift of 145 nm. ASCP has a higher binding affinity to fibrillar α-synuclein than ThT and likely shares the same binding sites to amyloid fibrils. Importantly, ASCP was found to also be fluorogenic in the presence of amorphous aggregates and can detect oligomeric species formed early during aggregation. Moreover, ASCP can be used to visualise fibrils via Total Internal Reflection Fluorescence (TIRF) microscopy and, due to its large Stokes shift, simultaneously monitor the fluorescence emission of other labelled proteins following excitation with the same laser used to excite ASCP. Consequently, ASCP possesses enhanced and unique spectral characteristics compared to ThT that make it a promising alternative for the *in vitro* study of amyloid fibrils and the mechanisms by which they form.

## Introduction

The aggregation of proteins into amyloid fibrils is associated with a variety of protein conformational disorders, including neurodegenerative diseases such as Alzheimer’s disease, Parkinson’s disease and Amyotrophic Lateral Sclerosis. Amyloid fibril formation typically occurs via a nucleation-dependent process ^1^, which results in the formation of soluble oligomeric species to which partially folded or unfolded monomers are added in a two-dimensional process to form a fibril ^2^. Amyloid fibrils are characterised by a well-defined cross β-sheet structure that runs perpendicular to the long axis of the fibrils ^3^. Much of the research into protein aggregation over the last few decades has focused on fibrillogenesis due to its close association with disease states ^4^. Whilst the toxicity of amyloid fibril formation has been recognised for some time, there has been considerable debate regarding which species is responsible for the observed cytotoxicity of amyloid fibrils. Increasing evidence suggests that soluble pre-amyloid oligomeric species, rather than the mature fibrils themselves, are the most cytotoxic species and may be responsible for disease pathogenesis ^5,6^. Consequently, there is a continuing need to develop *in vitro* approaches that enable the process of fibril formation to be better understood at the molecular level and in the context of disease onset and progression.

Amyloid-binding dyes have been extensively used to study the aggregation of proteins into amyloid fibrils *in vitro*. Thioflavin T (ThT) is the best characterised and the most commonly used fluorescent dye for the detection of amyloid fibrils ^7^. Structurally, ThT consists of a dimethylaminobenzene ring coupled to a positively charged benzothiazole ring by a rotatable C-C bond; internal rotation around this bond quenches the locally excited state of the molecule to a dark, twisted intramolecular charge transfer state that results in a low fluorescence yield of ThT when free in solution ^8^. When the intramolecular rotations of ThT are inhibited, such as upon binding to the β-sheet rich motifs that are enriched in amyloid, the excited state is preserved and the quantum yield of fluorescence increases dramatically ^9^. Unfortunately, ThT has a limited ability to detect oligomeric species that are present early during the aggregation into amyloid fibrils ^10,11^. Furthermore, ThT also has a relatively small Stokes shift (∼ 50 nm), which can be problematic when used in conjunction with other spectrally similar molecules ^12^. This is particularly relevant in the context of fluorescence imaging techniques that exploit the selective amyloid binding properties of ThT, such as Total Internal Reflection Fluorescence (TIRF) microscopy. Such methods have enabled the visualisation of individual amyloid fibrils ^13^, and facilitated the observation of fibril elongation in real-time ^14–18^. However, due to the relatively small Stokes shift of ThT, experiments that also require fluorescently labelled proteins can be technically challenging owing to the requirement of multiple excitation sources to avoid spectral overlap.

Recently, efforts have been made to develop novel aggregation-induced emission (AIE) fluorophores that have enhanced spectral qualities and can overcome some of the limitations of ThT ^19,20^. For example, a tetraphenylethene tethered to triphenylphosphonium (TPE-TPP) dye has been recently described with superior ability to detect oligomeric species formed early during the aggregation of proteins into amyloid fibrils ^21–23^. However, the maximum fluorescence emission for both ThT and TPE-TPP is in the blue region of the spectrum (i.e. an emission maximum of ∼ 480 nm) and this can restrict their use in cells or tissues due to significant interference as a result of autofluorescence from these samples. The present study aimed to characterise the AIE fluorophore ASCP for its suitability as a fluorescent probe to monitor amyloid fibril formation and the imaging of fibrils via TIRF microscopy. Whilst the design, synthesis and application of ASCP for imaging of cellular organelles has been previously described ^24^, here we show that ASCP is fluorogenic in the presence of species formed during the fibrillation and amorphous aggregation of proteins. Moreover, we demonstrate that ASCP has enhanced affinity to amyloid fibrils compared to ThT and has advantages over ThT in its use in single-molecule TIRF microscopy experiments as a result of its large Stokes shift that enables a single excitation source to be used to simultaneously monitor the emission from a fluorescently labelled protein and ASCP-stained aggregates.

## Materials and Methods

ThT was purchased from Sigma Aldrich while the ASCP dye was synthesised following previous reported procedures ^24^. Both dyes were solubilised in 100% DMSO to a final concentration of 5 mM. All buffers and solutions were filtered using 0.22 µm filters before use and contained 0.02% (w/v) NaN_3_ to prevent bacterial contamination. κ-casein, insulin and *α*-lactalbumin were obtained from Sigma Aldrich and Aβ_1-42_ was purchased from AnaSpec. κ-casein was reduced and carboxymethylated (RCM) as previously described to generate RCM κ-casein ^25^. Recombinant *α*-synuclein was expressed and purified as described ^26^. Recombinant *α*B-crystallin (*α*B-c) was expressed and purified as described previously ^27^, and fluorescently labelled with Alexa Fluor 488-maleimide (AF488) as per the manufacturers instruction to generate AF488-*α*B-c. AF488-labelled SOD1 was a gift from Prof. Justin Yerbury and Dr. Luke McAlary. All protein stocks were stored in aliquots at −20 °C until required.

### Aggregation of *α*-synuclein

Fibrils of *α*-synuclein were generated as described previously ^28^, with some modifications. Briefly, to produce α-synuclein seeds (fibril fragments), aliquots of monomeric α-synuclein (200 μM) were prepared in 20 mM phosphate buffer (pH 6.3) and incubated at 45°C with maximal stirring, using a Teflon flea on a WiseStir heat plate for 24 hr to facilitate the formation of fibrils. The sample was then sonicated using a Digital Sonifier 250 (Branson, USA) for three cycles of 10 sec at 30% power. The incubation and sonication steps were then repeated. The seed fibrils were collected via centrifugation at 50,000 x *g* for 20 min at 4°C and the resulting pellet containing the seed fibrils resuspended in 20 mM phosphate buffer (pH 7.4). Aliquots of the *α*-synuclein seeds were stored at -20°C. To produce mature α-synuclein fibrils, α-synuclein seeds (2.5 μM) were incubated with monomeric α-synuclein (50 μM) in 20 mM phosphate buffer (pH 7.4) at 45 °C without agitation for 48 hr. The fibrils were then centrifuged at 50,000 x g for 20 min at 4°C and the pellet containing the mature fibrils was resuspended in 20 mM phosphate buffer (pH 7.4) and stored at room temperature. The concentration of fibrils and seed fibrils are reported as the equivalent monomeric concentration used to prepare them.

### Absorbance and fluorescence spectra of dyes

Absorbance spectra of ThT and ASCP dyes were determined on a Nanodrop 2000C (ThermoFisher). Spectra were determined in 20 mM phosphate buffer (pH 7.4) and in the presence of monomeric and fibrillar *α*-synuclein (50 µM) to determine if any shift in absorbance occurs upon binding of dyes to the protein. Spectra were generated with 100 µM of dye. Data was corrected for signals produced by an equivalent concentration of DMSO alone.

The fluorescence spectrum of ThT and ASCP was determined using a Varian Cary Eclipse Fluorometer, following excitation at the maximum absorbance wavelength for the individual dye (412 or 460 nm for ThT and ASCP, respectively). The fluorometer was set with excitation and emission slit widths of 5 nm and 10 nm, respectively, and data collected at a scan rate of 300 nm/min and an averaging time of 0.5 sec. All samples were loaded into a quartz fluorescence cuvette. Increasing concentrations of each dye (0 - 50 µM) in the presence of monomeric or fibrillar *α*-synuclein (20 µM) were tested. Controls consisted of dye alone (20 µM) without *α*-synuclein, and *α*-synuclein (20 µM, monomer or fibril) in the presence of an equivalent concentration of DMSO alone. Values from appropriate controls were subtracted from the corresponding raw data.

### Binding affinity and binding competition of ThT and ASCP to α-synuclein fibrils

To determine the binding affinity of ThT and ASCP to *α*-synuclein fibrils, the fluorescence intensity at the maximum emission wavelength (480 nm for ThT and 605 nm for ASCP) for each concentration of dye in the presence of *α*-synuclein fibrils was determined and normalised against the maximum fluorescence value. Each normalised peak fluorescence was plotted against dye concentration and the apparent equilibrium binding constant (*K*_*b*_) was determined by fitting the data to a one-phase association model using GraphPad Prism8 software.

To determine whether ASCP can out-compete ThT for binding sites on fibrillar *α*-synuclein, ThT (20 μM) was incubated in the absence or presence of *α*-synuclein fibrils (20 μM) in 20 mM phosphate buffer (pH 7.4) at room temperature for 5 min. Different concentrations of ASCP (0 - 20 μM) were then added to *α*-synuclein samples containing ThT and the fluorescence intensities for ThT (440 nm excitation, 490 emission) and ASCP (485 nm excitation, 620 nm emission) were recorded using a FLUOStar plate reader (BMG Labtech). The fluorescence values were normalised to the maximum fluorescence intensity, plotted against ASCP concentration and the data was fit to a one-phase association model using GraphPad Prism8 software.

### Monitoring the fibrillar and amorphous aggregation of RCM K-casein, Aβ_1-42_ peptide, *α*-lactalbumin and insulin using ThT and ASCP *in situ*

To test the ability of ASCP to monitor the formation of amyloid fibrils by proteins other than *α*-synuclein, ThT (20 µM) or ASCP (2 - 20 µM) was added to either RCM *κ-*casein (50 µM) in 50 mM phosphate buffer (pH 7.4) or Aβ_1-42_ peptide (20 µM) in phosphate buffered saline (PBS). To test whether ASCP was selectively fluorogenic in the presence of amyloid aggregates, the amorphous aggregation of *α*-lactalbumin and insulin in the presence of ThT and ASCP was also monitored. *α*-lactalbumin in 50 mM imidazole (pH 7.0) supplemented with 5 mM CaCl_2_ and 100 mM NaCl or insulin in 50 mM phosphate buffer (pH 7.4) were incubated in the presence of various concentrations of dye (2 – 20 µM). Aggregation of *α*-lactalbumin and insulin was initiated following addition of dithiothreitol (DTT, 20 mM or 2 mM, respectively).

Aggregation assays were performed in a FLUOstar or POLARstar plate reader set at either 37°C for RCM κ-casein, *α*-lactalbumin and insulin or 30°C for Aβ_1-42_ peptide. For amyloid forming proteins, assays were performed in black 384 well plates and the fluorescence was measured every 12 min (RCM κ-casein) or 24 min (Aβ_1-42_) until fluorescence signal had plateaued (∼ 48 or 72 hr for RCM κ-casein or Aβ_1-42_, respectively). For *α*-lactalbumin and insulin, assays were performed in black 96 well plates and the absorbance at 340 nm and the fluorescence of the dyes was measured alternatively every 3.3 min for 300 min total. For the aggregation of RCM κ-casein, Aβ_1-42_ and insulin, plates were shaken for 30 sec before each cycle. The following excitation/emission wavelengths were used; ThT *λ*_ex_ = 440 nm, *λ*_em_ = 490 nm; ASCP *λ*_ex_ = 485 nm, *λ*_em_ = 590 nm. The change in fluorescent intensity is reported for each dye, while results were normalised by dividing the fluorescence intensity at each time point by the maximum fluorescence for easy comparison of dyes. RCM κ-casein assays were run in duplicate, while Aβ_1-42_, *α*-lactalbumin and insulin aggregation assays were run in triplicate. Dye alone (20 µM) in the appropriate assay buffer served as a control for each sample. Data from the RCM κ-casein assay were fitted with a specific binding with hill slope model using GraphPad Prism8 software.

### ThT and ASCP as *in situ* reporters of α-synuclein fibril elongation

The seeded aggregation of α-synuclein was monitored *in situ* using a microplate assay, as previously described ^29^. Duplicate samples, containing varying concentrations of ThT or ASCP (0 - 50 μM) in the presence or absence of α-synuclein (50 μM monomeric, 2.5 μM seed) were incubated in 20 mM phosphate buffer (pH 7.4) for 48 hr at 37°C without shaking. The fluorescence was measured every 15 min using the excitation and emission filters of 440/490 nm for ThT and 485/610 nm for ASCP. The fluorescence values from samples in the absence of α-synuclein were subtracted from equivalent readings containing α-synuclein to correct for signals produced by the dye alone (which were negligible). The fluorescence intensity values obtained from each sample of the assay were normalised to the maximum fluorescence intensity of that sample. The raw ASCP fluorescence values and the normalised values from both dyes were plotted as a function of time and fitted with a one-phase association model using GraphPad Prism8 software.

At the end of the *in situ* α-synuclein fibril formation assay an aliquot was taken from each sample and centrifuged at 50,000 x g for 20 min at 4°C to separate the supernatant, which contains soluble monomeric α-synuclein, from the pellet that contains the fibrillar α-synuclein. The supernatants were collected, and the pellets were resuspended to the same initial volume using 20 mM phosphate buffer (pH 7.4) for subsequent analysis via SDS-PAGE.

### Assessing the ability of ThT and ASCP to monitor the aggregation of *α*-synuclein *ex situ*

*α*-synuclein was aggregated as described and aliquots of *α*-synuclein were taken at *t =* 0, 1, 3, 6, 12, 24 and 72 hr following the start of aggregation and stored at 4°C to prevent further aggregation. The fluorescence spectra of *α*-synuclein fibrils (20 µM) in the presence of dye (20 µM) at each time point was collected using a fluorometer set with excitation and emission slit widths of 5 nm and data acquired at a scan rate of 200 nm/min and an averaging time of 0.2 sec. ThT and ASCP were excited with their previously determined maximum absorbance wavelengths (412 or 460 nm for ThT and ASCP, respectively). The maximum fluorescence value for each timepoint was determined and normalised to the maximum fluorescence intensity of the 72 hr timepoint. The normalised fluorescence values were graphed and fit with a one-phase association curve using GraphPad Prism8 software.

### Preparing coverslips for TIRF microscopy

To prepare slides for microscopy, 24 x 24 mm glass coverslips were cleaned by alternatively sonicating for 30 min in 100% ethanol then in KOH (2 M) for a total of 2 hr, and with a final sonication in milli-Q water for 5 min. The coverslips were then incubated for 30 min at room temperature with poly-L-lysine (0.01%, w/v) and washed three times with 20 mM phosphate buffer (pH 7.4). Coverslips, with loaded sample, were placed onto the TIRF objective lens coated with immersion oil (n = 1.518, Olympus, UK).

### TIRF microscopy instrument setup

Samples were imaged using a home-built inverted optical microscope (Nikon Eclipse TI) that was coupled to an electron multiplied charged coupled device (EMCDD) camera (Evolve II 512, Photometrics, Tuscon, AZ). The camera was established to operate in an objective type TIRF setup with a diode-pumped solid-state laser (200 mW Sapphire, Coherent, USA) emitting a circularly polarized laser radiation of 488 nm continuous wavelength. The emitted light was directed off a dichroic mirror (ZT405/488/561/647 Semrock) through an oil immersion objective lens (CFI Apochromate TIRF Series 60x objective lens, numerical aperture = 1.49) and onto the sample. Total internal fluorescence was achieved by directing the incident ray onto the sample at an angle greater than the critical angle of θ_c_ ∼ 67**°** for a glass/water interface. The evanescent light field generated selectively excites the fluorophores, with the emission passing through the same objective lens and filtered by the same dichroic mirror. The emission was then passed through a 635 nm long pass filter (BLP01-635R, Semrock, USA) and the final fluorescent image projected onto the EMCDD camera. The camera was running in frame transfer mode at 20 Hz, with an electron multiplication gain of 700, operating at - 70°C with a pixel.

### Monitoring α-synuclein fibril elongation *ex situ* using TIRF microscopy

The seeded aggregation of α-synuclein was monitored *ex situ* using TIRF microscopy. Fibrillar *α*-synuclein was generated as described above, with the exception that samples were incubated at 37°C, and aliquots were taken at various time points (0, 1, 3, 6, 12, and 24 hr). Aliquots were centrifuged at 50,000 × g for 20 min at 4°C to remove monomeric α-synuclein that was not incorporated into the fibrils. Samples were diluted 4,000-fold into 20 mM phosphate buffer (pH 7.4) containing 6 mM 6- hydroxy- 2,5,7,8-tetramethylchroman-2-carboxylic acid (trolox) and ASCP (5 µM). The sample (50 μL) was then loaded onto the coverslip and imaged. Prior to imaging, samples were randomised so the user was blinded to the sample identity.

### Direct observation of αB-crystallin (*α*B-c) binding to fibrillar *α*-synuclein

To demonstrate the ability of ASCP to be used in single-laser two-colour TIRF microscopy experiments, the well-characterised interaction between the small heat-shock protein (sHsp) *α*B-crystallin (αB-c, HspB5) and mature α-synuclein fibrils was investigated ^29^. Samples containing α-synuclein fibrils (50 μM) and AF488-labelled αB-c (1 μM) were incubated together at room temperature for 5 min in 20 mM phosphate buffer (pH 7.4) containing 6 mM trolox and diluted 1,000-fold into the same buffer containing ASCP (5 μM). Samples (50 μL) were loaded onto the coverslip and imaged. Samples were randomised prior to imaging so the user was blinded to the sample identity.

### Image analysis

Using custom plugin software operated through Fiji ^30^, background fluorescence and laser beam profiles were subtracted from the acquired images. The images were Z-projected, averaging all the frames attained during acquisition, and the brightness and contrast adjusted for clear visualisation.

## Results

### Absorbance and fluorescence spectra of ThT and ASCP

Absorption spectra were acquired to determine the maximum absorbance wavelength (*λ*_max_) and the effect of monomeric and fibrillar *α*-synuclein on the absorbance of ThT and ASCP (Figure 1A and B). The maximum absorbance for ThT and ASCP in solvent alone or in the presence of monomeric *α*-synuclein was 412 nm and 430 nm, respectively. There was no shift in *λ*_max_ for ThT when bound to fibrillar *α*-synuclein. Interestingly, ASCP was observed to have a higher absorbance in the presence of monomeric or fibrillar *α*-synuclein and exhibited a significant shift (30 nm) in *λ*_max_ to 460 nm when bound to fibrillar *α*-synuclein. Based on the absorbance spectra, excitation wavelengths (*λ*_ex_) of 412 nm (ThT) and 460 nm (ASCP) were used to examine the fluorescent properties of the dyes in solvent alone or bound to *α*-synuclein.

**Figure 1:**
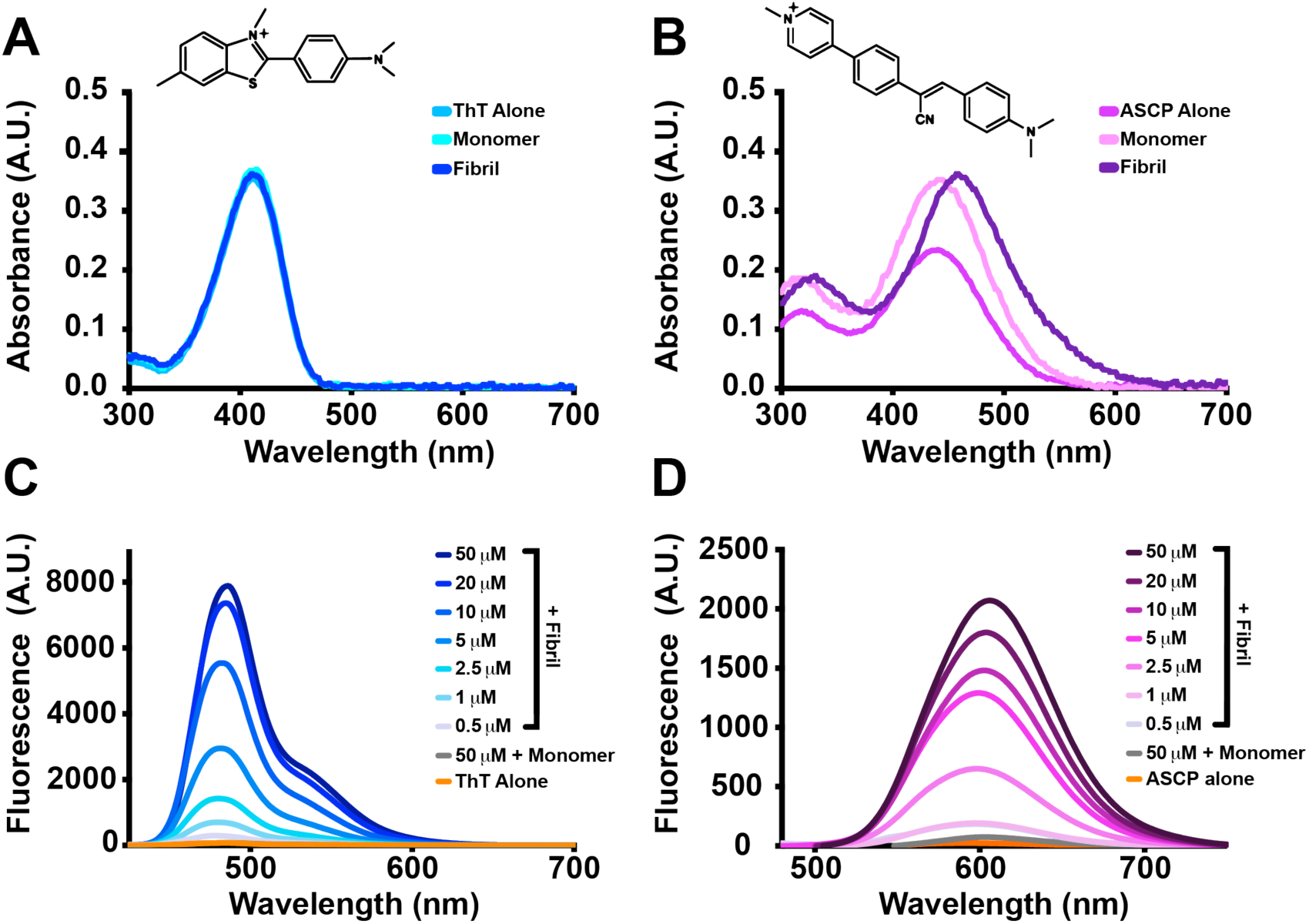
Absorbance and fluorescence spectra of ThT and ASCP in the presence or absence of monomeric or fibrillar *α*-synuclein. The dyes ThT **(A)** and ASCP **(B)** (100 µM) were added to 20 mM phosphate buffer (pH 7.4) in the presence or absence of monomeric or fibrillar *α*-synuclein (50 µM). Absorbance spectra was measured from 300 - 700 nm. The chemical structures of ThT (A, *inset*) and ASCP (B, *inset*) are shown. The fluorescence emission spectra of both ThT **(C)** and ASCP **(D)** (0 - 50 µM) in the presence of monomeric or fibrillar *α*-synuclein (20 µM) was then measured using a spectrofluorometer, following excitation at 412 nm and 460 nm for ThT and ASCP, respectively.

Both ThT and ASCP demonstrated significant fluorescent enhancement when bound to fibrillar *α*-synuclein as compared to when the dye was in solvent alone or in the presence of monomeric *α*-synuclein (Figure 1C and D). The fluorescence intensity of 50 µM of ThT and ASCP was minimal when in solvent alone with negligible increases in fluorescent intensity when bound to monomeric *α*-synuclein. The maximum emission wavelength (*λ*_em_) for ThT (50 µM) was 480 nm, while ASCP (50 µM) exhibited a significant Stokes shift of ∼ 145 nm with a *λ*_em_ of 605 nm.

### Binding affinity and binding competition of ThT and ASCP to *α*-synuclein fibrils

The normalised fluorescence at each concentration of ThT and ASCP was used to determine the apparent binding constant (*K*_*b*_) of each dye to fibrillar *α*-synuclein (Figure 2A). By fitting a one-phase association model, the binding affinity of ASCP (*K*_*b*_ = 5.5 ± 0.7 µM) was determined to be almost twice that of ThT (*K*_*b*_ = 9.9 ± 1.5 µM). Competition experiments were then performed to determine whether ASCP outcompetes ThT for binding to fibrillar *α*-synuclein (Figure 2B). In the presence of *α*-synuclein fibrils that had been pre-incubated with ThT (20 μM), increasing concentrations of ASCP resulted in an increase in the intensity of ASCP fluorescence and a simultaneous decrease in ThT fluorescence intensity. At a molar ratio of 1:2 (ASCP:ThT) there was a > 95% reduction in ThT fluorescence, which indicated that ThT was competitively dissociated from fibrils by ASCP.

**Figure 2:**
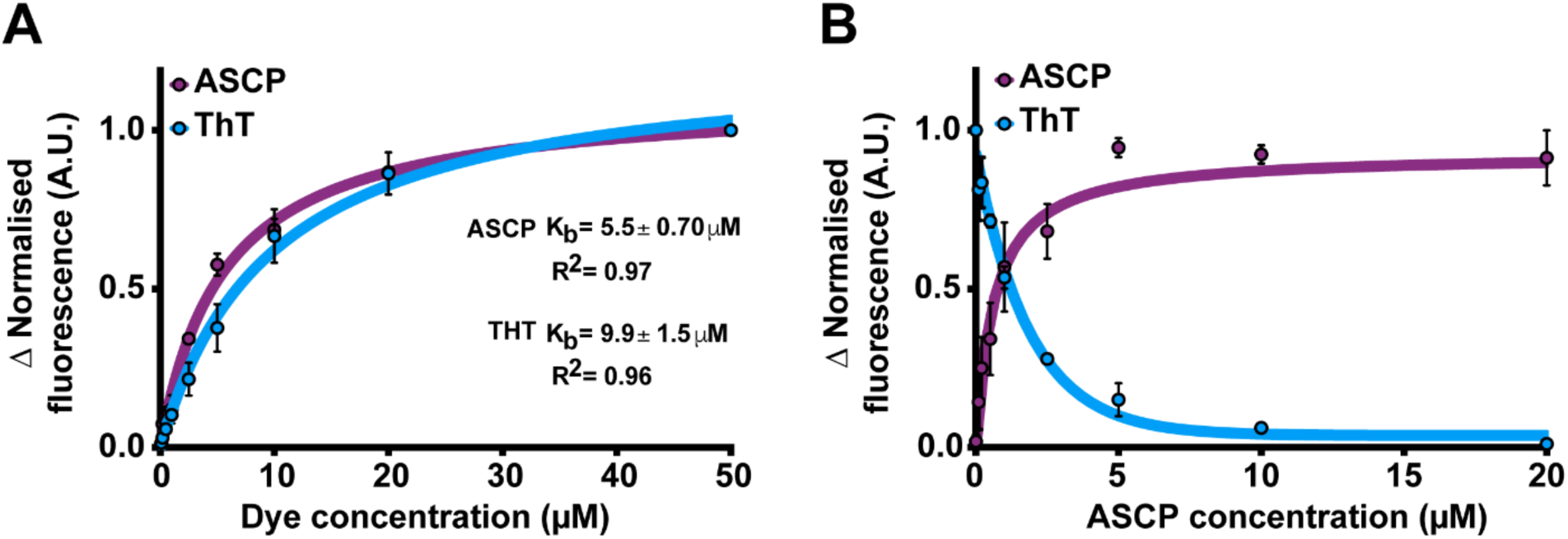
Binding affinity and competitive binding of ASCP and ThT to *α*-synuclein fibrils. **(A)** ThT or ASCP (0 – 50 µM) was incubated with *α*-synuclein fibrils (20 µM). The normalised fluorescence intensity was plotted against the dye concentration and fitted to a one-phase association model. The calculated apparent binding affinity (*K*_*b*_) is shown for each dye. **(B)** ASCP (0 - 20 µM) was added to *α*-synuclein fibrils (20 µM) pre-incubated in the presence of ThT (20 µM). The normalised fluorescence intensity of each dye was plotted against the concentration of ASCP. The data corresponding to ASCP fluorescence was fitted to a one-phase association model and ThT fluorescence fitted to a one-phase dissociation model. Data represents the mean ± standard error from three independent experiments.

### Monitoring the fibrillar and amorphous aggregation of various model proteins *in situ*

In order to determine whether ASCP is capable of monitoring the aggregation of other proteins into amyloid fibrils, Aβ_1-42_ peptide and RCM κ-casein aggregation assays were performed (Figure 3). The fluorescence intensity of both ThT and ASCP (20 µM) were observed to increase over time when incubated with either Aβ_1-42_ or RCM κ-casein. Interestingly, the fluorescence intensity profile of ASCP differed substantially to ThT during the aggregation of Aβ_1-42_, with ASCP fluorescence increasing early during the aggregation process (0 – 40 hr) before plateauing after 40 hr (Figure 3A). Conversely, the fluorescence intensity of ThT only increased substantially after 40 hr of aggregation and plateaued after ∼ 55 hr (Figure 3A). When used to monitor the aggregation of Aβ_1-42_ the fluorescence intensity profile of ASCP did not change between concentrations of 2 to 20 μM (Figure 3A), suggesting that the dye itself does not significantly influence the kinetics of Aβ_1-42_ aggregation. Similar as for Aβ_1-42_, when ASCP and ThT were used to monitor amyloid fibril formation by RCM κ-casein the increase in ASCP fluorescence occurred much earlier than for ThT (Figure 3B), with the time to half maximum intensity determined to be 8.6 hr for ASCP and 15.6 hr for ThT. These data again suggest that ASCP may detect species formed earlier during the aggregation of RCM κ-casein into amyloid fibrils than ThT.

**Figure 3:**
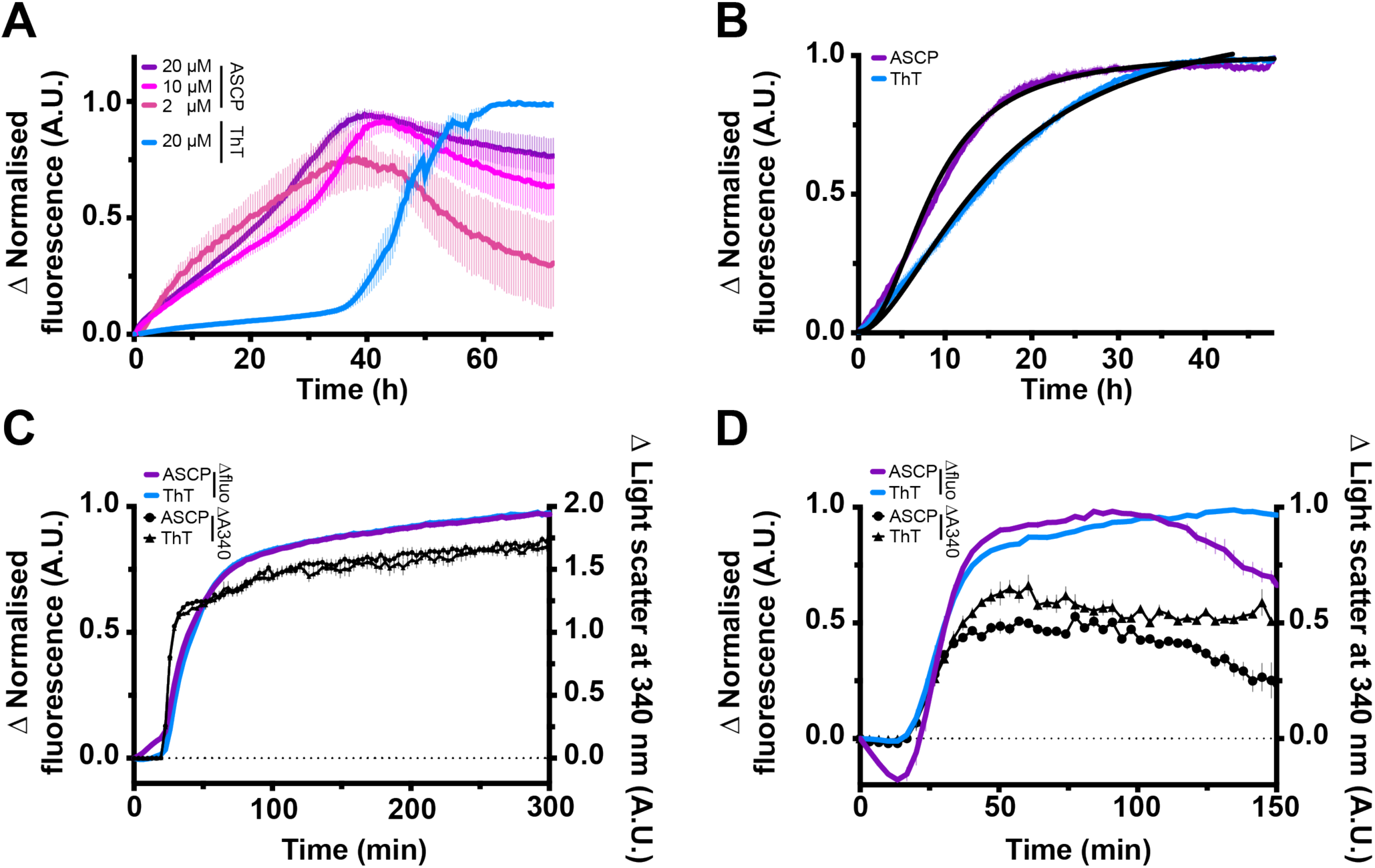
ASCP can be used to monitor the fibrillar and amorphous aggregation of proteins *in situ.* ThT or ASCP (20 µM) were added to Aβ_1-42_ (20 µM, **A**), RCM κ-casein (50 µM, **B**), *α*-lactalbumin (100 µM, **C**) or insulin (100 µM, **D**) and the aggregation of protein was monitored via fluorescence and/or light scatter at 340 nm. For monitoring the aggregation of Aβ_1-42_, various concentrations of ASCP (2 – 20 µM) were present during the incubation. The fluorescence was measured using excitation/emission wavelengths of 440/490 nm and 485/590 nm for ThT and ASCP, respectively. RCM κ-casein assays were run in duplicate, while Aβ_1-42_, *α*-lactalbumin and insulin aggregation assays were run in triplicate. Data for all panels is shown as the mean ± standard error from at least 3 independent experiments.

To determine whether ASCP could monitor the amorphous aggregation of proteins, the reduction-induced aggregation of *α*-lactalbumin and insulin was monitored in the presence of ThT and ASCP (Figure 3C and D). An initial lag phase of ∼ 20 min was observed for both *α*-lactalbumin and insulin, whereby there was no increase in light scatter at 340 nm during this period. After the observed lag phase, the light scatter at 340 nm was observed to increase for both *α*-lactalbumin and insulin, indicative of the formation of amorphous aggregates. The change in light scatter was observed to plateau after ∼ 120 min and 50 min for *α*-lactalbumin and insulin, respectively. Interestingly, the fluorescence intensity of both ThT and ASCP also increased in these samples in a similar manner to that observed for the increase in light scatter at 340 nm. Notably, there was an increase in fluorescence intensity observed for ASCP, but not ThT, during the early stages (0 – 20 min) of *α*-lactalbumin aggregation, a time period corresponding to the observed lag phase as determined by light scattering at 340 nm. There was no difference in the magnitude or profile of the change in light scatter at 340 nm when *α*-lactalbumin or insulin were incubated in the absence or presence of the dyes (Supplementary Figure 1), suggesting that the dyes themselves do not significantly influence the aggregation kinetics of these two proteins.

### ASCP and ThT can be used *in situ* to monitor the aggregation of *α*-synuclein in real time

*In vitro* aggregation assays were used to compare the abilities of ASCP and ThT to monitor *α*-synuclein fibril elongation *in situ* over time. The addition of *α*-synuclein seeds removes the lag phase kinetics associated with *α*-synuclein aggregation ^28^, and therefore enables fibril elongation to be examined. When *α*-synuclein was incubated in the presence of ThT, there was an increase in the normalised fluorescence intensity at 490 nm over time that corresponds to the elongation of *α*-synuclein fibrils (Figure 4A). At all concentrations of ThT tested, there was a uniform increase in fluorescence intensity and a plateau observed after ∼ 24 hr of incubation. Similarly, there was an increase in the normalised fluorescence intensity at 610 nm observed when *α*-synuclein was incubated in the presence of ASCP (Figure 4B). A similar plateau in fluorescence intensity was also reached following 24 hr of incubation in the presence of up to 5 μM ASCP; however, at concentrations above 5 μM of ASCP a plateau was not yet reached after 48 hr of incubation and the initial rate of elongation was reduced. The total change in ASCP fluorescence intensity at the endpoint of the assay, which can provide an indication on the total amount of amyloid formed, was observed to increase in magnitude up to 10 µM of ASCP and decrease at concentrations greater than 10 µM (Figure 4C).

**Figure 4:**
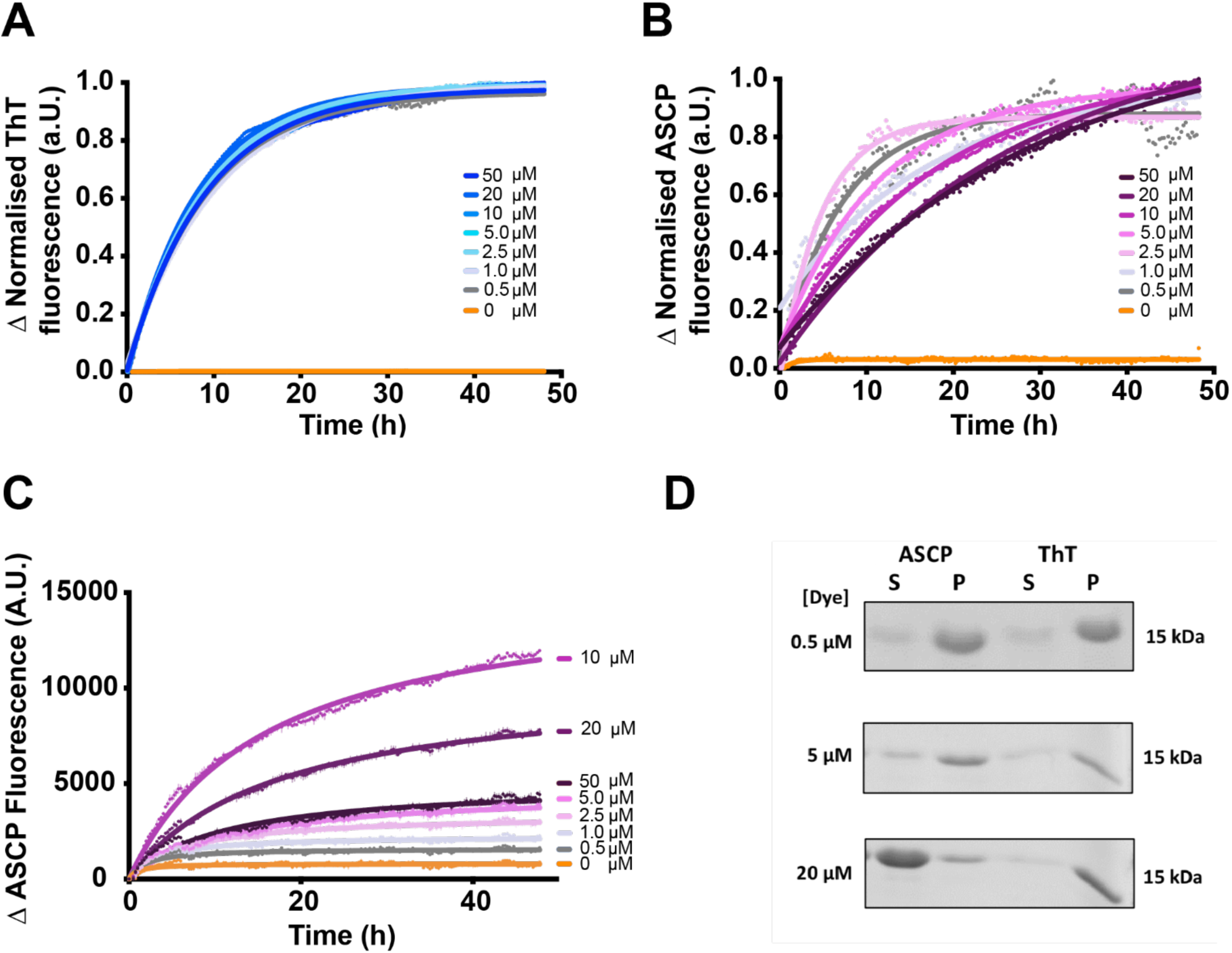
The *in situ* elongation of α-synuclein fibrils measured via ThT or ASCP fluorescence. Monomeric α-synuclein (50 μM) and α-synuclein seeds (2.5 μM) in 20 mM phosphate buffer (pH 7.4) was incubated at 37°C in the presence of varying concentrations of dye (0 – 50 µM). The elongation of the seeds was monitored by measuring the change in fluorescence over time of ThT (**A**) or ASCP (**B**), which was normalised to allow for easy comparison between the dyes. (**C**) The relative change in fluorescence intensity during the elongation of *α*-synuclein seeds in the presence of different concentrations of ASCP (0 – 50 µM). The data in A-C are representative of 3 independent repeats. **(D)** Samples from the microplate assays (from panels A and B) were centrifuged at 50,000 x g for 20 min to separate soluble monomeric α-synuclein fractions ‘S’ from α-synuclein fibrils within the pellet ‘P’ and subsequently analysed via SDS-PAGE.

To establish whether the reduced fluorescence intensity and rates of *α*-synuclein fibril elongation at higher concentrations of ASCP is due to inhibition of fibril elongation, samples were collected at the end of the assay and the proportion of soluble (supernatant) and fibrillar (pellet) *α*-synuclein was analysed via SDS-PAGE (Figure 4D). When incubated in the presence of ThT, *α*-synuclein (denoted by a band at ∼ 15 kDa) was primarily observed in the insoluble pelleted fraction, indicating that elongation of *α*-synuclein seeds into fibrils had occurred in the presence of all concentrations of ThT tested. Similar to ThT, and in agreement with the elongation assays, *α*-synuclein was observed predominantly in the insoluble pelleted fractions when incubated in the presence of up to 5 µM ASCP. However, at concentrations of ASCP greater than 5 μM, *α*-synuclein was found predominantly in the soluble (supernatant) fraction. Thus, these data suggest that high concentrations of ASCP inhibits *α*-synuclein fibril elongation.

### Monitoring and visualising the aggregation of *α*-synuclein using *ex situ* assays and TIRF microscopy

To determine if ASCP could be used to monitor the seeded aggregation of *α*-synuclein *ex situ, α*-synuclein was incubated in the absence of dye and aliquots taken at various time intervals and incubated with ThT or ASCP. The normalised fluorescence of both ThT and ASCP in the presence of *α*-synuclein was observed to increase over time (Figure 5A), indicative of the formation of amyloid fibrils. The increase in fluorescence of both ThT and ASCP over time fits well to a one-phase association curve, which is typical for the seeded aggregation of *α*-synuclein ^28^.

**Figure 5:**
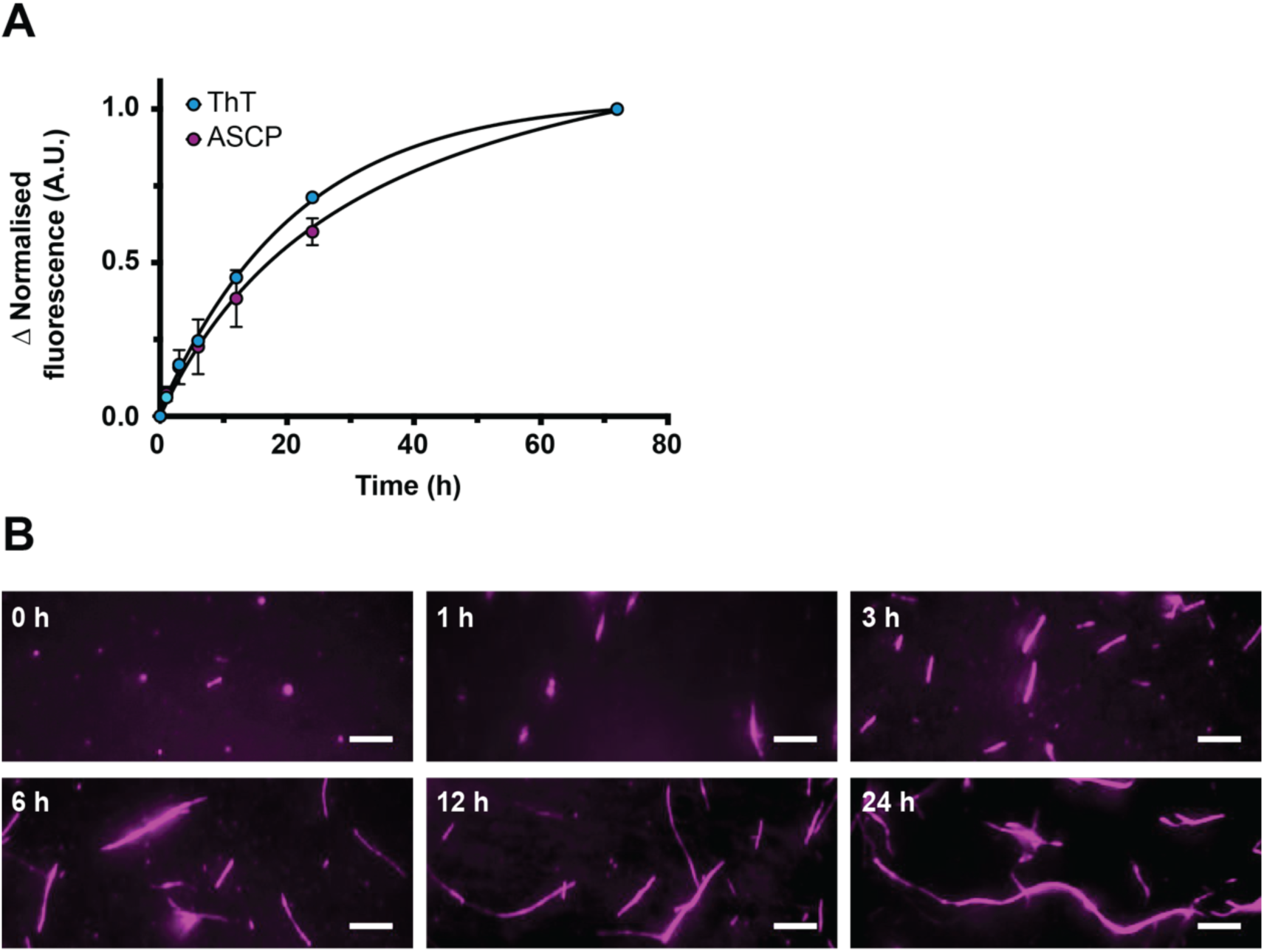
Monitoring and visualising the elongation of *α*-synuclein fibrils using *ex situ* fluorometry and TIRF microscopy. Recombinant *α*-synuclein monomer (50 µM) was incubated in the presence of *α*-synuclein seeds (2.5 µM) in 20 mM phosphate buffer (pH 7.4) to generate mature fibrils. **(A)** Aliquots were taken from the elongation reaction at 0, 1, 3, 6, 12 and 24 and 72 hr. The generated *α*-synuclein fibrils (20 µM final concentration) at each time point were incubated in the presence of either ThT or ASCP (20 µM) and the fluorescence measured. The normalised fluorescence intensity was plotted against time and fitted to a one-phase association model. **(B)** Aliquots were taken from the elongation reaction at 0, 1, 3, 6, 12 and 24 hr. Samples from each timepoint were diluted 4,000-fold in 20 mM phosphate-trolox buffer containing ASCP (5 µM) and imaged using TIRF microscopy. Representative TIRF microscopy images from each timepoint are shown. Scale bar represents 5 µm.

As it was established that ASCP could be used to monitor the elongation of α-synuclein fibrils *in situ and ex situ*, we next examined whether the dye could be used to visualise fibrils using TIRF microscopy (Figure 5B). To do so, aliquots from a seeded aggregation reaction were taken at various timepoints, incubated with ASCP and deposited onto a Poly-L-lysine coated coverslip for image acquisition. Fibrils could clearly be visualised via TIRF microscopy following excitation of 488 nm and using an emission window of > 675 nm. By using ASCP and TIRF microscopy to visualise individual fibrils, the length of the α-synuclein fibrils was observed to increase during incubation; short fluorescent fragments corresponding to α-synuclein seeds were visible at 0 hr and thin filament-like structures, typical of amyloid fibrils and with some exceeding lengths of 10 µm, were observed over 24 hr. Importantly, ASCP displayed negligible fluorescence in the presence of monomeric α-synuclein; however, a small number of fluorescent puncta were observed in some fields of view, typical of insoluble dye molecules (Supplementary Figure 2).

After establishing that ASCP can be used to visualise *α*-synuclein fibrils using TIRF microscopy, we sought to further exploit the unique spectral properties of ASCP over ThT, in particular the large Stokes shift in its emission, by performing single-laser two-colour microscopy. As a proof of principle, the ability of the molecular chaperone *α*B-crystallin (*α*B-c) to bind to *α*-synuclein fibrils was investigated. As part of its chaperone function, αB-c inhibits α-synuclein fibril elongation, in part, through its ability to interact and form stable complexes with preformed, mature α-synuclein fibrils ^13,31–33^. Thus, mature *α*-synuclein fibrils were incubated in the presence of AF488-labelled *α*B-c (AF488-*α*B-c), diluted and imaged via TIRF microscopy using a single 488 nm laser (Figure 6). Upon excitation at 488 nm, the fluorescence emission of *α*-synuclein fibrils stained with ASCP was measured at > 675 nm while the fluorescence emission of AF488-*α*B-c bound to the fibrils was simultaneously measured at 500 - 550 nm. In the absence of *α*-synuclein fibrils, αB-c was found to be randomly dispersed on the coverslip surface and not co-localised to fluorescent puncta formed by insoluble ASCP, indicating that ASCP does not bind to αB-c (Supplementary Figure 2). However, when incubated in the presence of α-synuclein fibrils, αB-c was able to be visualised bound along the surface of the fibrils (Figure 6). In contrast, the non-chaperone control protein AF488-labelled SOD1 was found to be randomly dispersed in areas surrounding the α-synuclein fibrils, but not specifically associated with the fibrils (Supplementary Figure 2). Thus, using ASCP and TIRF microscopy we were able to simultaneously image *α*-synuclein fibrils and bound αB-c from the one laser light source.

**Figure 6:**
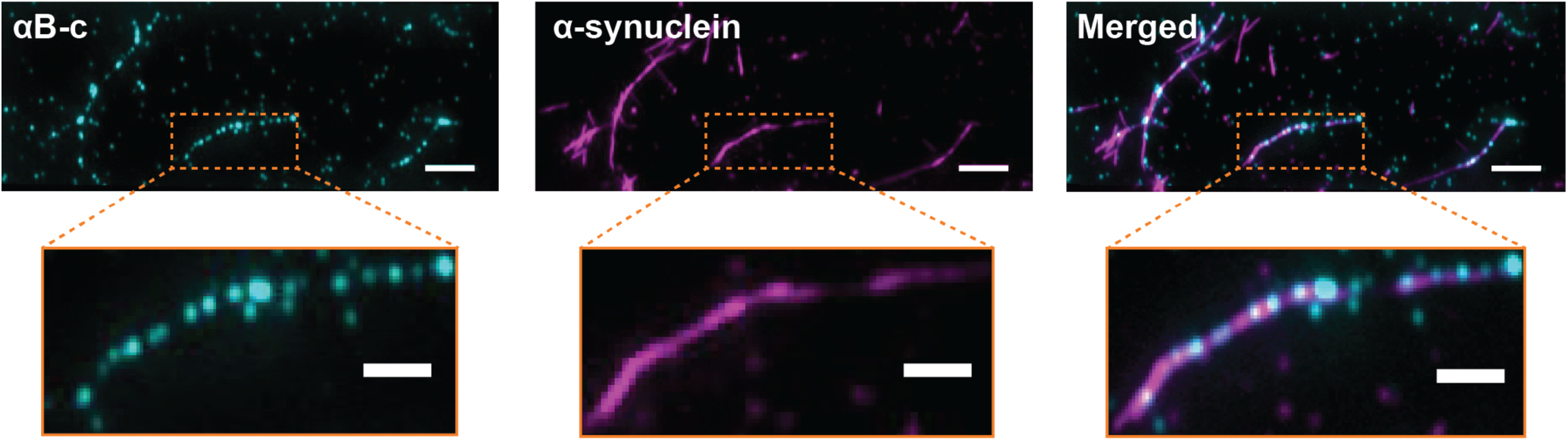
ASCP can be used in single-laser two-colour TIRF microscopy experiments to visualise binding of the molecular chaperone *α*B-c to *α*-synuclein fibrils. *α*-synuclein fibrils (50 µM) were incubated in 20 mM phosphate-trolox buffer (pH 7.4) in the presence of AF488-*α*B-c (1 µM) for 30 min at room temperature. Samples were diluted 1,000-fold in the presence of ASCP (5 µM final concentration) and both *α*-synuclein fibrils and AF488-*α*B-c was excited following illumination at 488 nm. A representative TIRF microscopy image of *α*B-c (*left, cyan*), *α*-synuclein stained with ASCP (*middle, magenta*) and a merged composite (*right*) is shown. Scale bars represent 5 µm or 2 µm (*magnified inset*).

## Discussion

Thioflavin T (ThT) is the most characterised and widely utilised fluorescent probe for the study and detection of amyloid fibrils. However, ThT is less able to detect oligomeric or pre-fibrillar species formed early during aggregation and possesses a small Stokes shift that complicates experiments that also employ spectrally similar fluorophores. Therefore, there is a need to identify and characterise other fluorogens with unique spectral properties that can be used to detect and monitor amyloid fibril formation. The aim of the present study was to characterise a red-emitting aggregation-induced emission (AIE) fluorogenic dye, ASCP, as a reporter of amyloid fibrils and assess its use for *in situ* and *ex situ* applications. Furthermore, we assessed the ability of ASCP to be used in TIRF microscopy-based experiments aimed at visualising and characterising individual *α*-synuclein amyloid fibrils and their interaction with molecular chaperones. Based on this work, we report that ASCP serves as an excellent alternative for ThT with regards to detecting and monitoring fibril formation and possesses the ability to detect species formed early during the aggregation of proteins into amyloid fibrils. Moreover, ASCP has spectral qualities that make it superior to ThT and can be used in single-laser TIRF-based investigations of amyloid fibrils.

The absorbance and fluorescence emission spectral properties obtained for both ThT and ASCP were similar to those previously reported ^11,24^. Whilst no changes were observed for the absorbance spectra of ThT in the presence or absence of α-synuclein fibrils, the absorbance spectra of ASCP increased in magnitude by ∼ 60% and was red-shifted 30 nm in the presence of α-synuclein. These changes in absorbance have been reported for other AIE dyes upon binding to amyloid structures ^34–36^, and has been specifically observed when ASCP is bound to DNA or RNA structures ^24^. The shift in ASCP absorbance spectra when in the presence of fibrillar α-synuclein is likely due to changes in local polarity caused by electrostatic solute-solvent interactions ^37,38^. Binding of the positively charged pyridinium moiety of ASCP to amyloid structures would result in a change of local polarity between the ground state (i.e. unbound and non-fluorescent) and excited state (i.e. bound and fluorescent) of the dye molecule. The resulting changes in polarity likely lead to the variations in absorbance magnitude and *λ*_abs_ observed for ASCP in the presence of different α-synuclein species.

Upon excitation at 460 nm (i.e. λ_ex_), ASCP displayed a maximum emission (λ_em_) at 605 nm, corresponding to a substantial Stokes shift of 145 nm, which is much greater than the Stokes shift exhibited by ThT (68 nm). Spectral shifts of this type can be attributed to changes in the intramolecular conformation and electronic structure of the dye as a result of the restriction of carbon-carbon bond rotations that occurs when the dye binds to amyloid structures ^39^. Restriction to a planar conformation results in the expansion of electron-π conjugates, (i.e. the adjacent donor-π-acceptor electron bridges of C=C bonds) and π-π stacking between benzene groups of dye molecules, all of which increase electron transfer and reduce the energy of the excited state ^40–42^. Thus, the enhanced Stokes shift of ASCP is attributed to the addition of a methylpyridinium and dimethylamine moieties to the *α*-cyanostilbene core of ASCP that enables an improved capacity to expand electron-π conjugates compared to that of ThT.

As demonstrated by both fluorometry and TIRF microscopy, negligible fluorescence was observed for ThT and ASCP in the presence of monomeric α-synuclein or solvent alone, whereas there was a substantial increase in fluorescence intensity in the presence of fibrillar α-synuclein. Both ThT and ASCP are known as fluorescent molecular rotors, with varying degrees of intramolecular rotation between C-C bonds. The rotation of C-C bonds quenches the locally excited states generated by photon excitation, resulting in a dark twisted intramolecular charge-transfer (TICT) state that produces little to no fluorescence emission ^43^. Restricting the rotational intermolecular motions of the dye, which is predicted to occur upon binding to amyloid fibrils ^8^, prevents energy transfer to the TICT state and preserves the excited state such that a high quantum yield of fluorescence occurs ^43^. There are at least three rotation sites within the structure of ASCP; either side adjacent to the C=C bond within its α-cyanostilbene backbone ^42^, and another in the C-C bond between the α-cyanostilbene backbone and methylpyridinium moiety. We propose that restriction of one or all of the rotational sites within ASCP results in the observed increase in fluorescence when bound to fibrillar α-synuclein.

The binding affinity of ASCP to α-synuclein fibrils was measured to be nearly twice that of ThT. This higher affinity is most likely due to additional electrostatic and hydrophobic interactions that can be made between ASCP and α-synuclein fibrils. Specifically, hydrophobic interactions between amyloid binding dyes and aromatic residues within fibrils are thought to facilitate high affinity interactions by enabling π-stacking and presenting large surfaces for dye binding ^9,44–46^. It is possible that the spatial separation and additional rotational permutations of the aromatic moieties of ASCP allow it to preferentially form hydrophobic contacts with the aromatic side chains of *α*-synuclein fibrils relative to ThT, contributing to the enhanced affinity of ASCP for amyloid observed in this study.

The simultaneous increase in fluorescence intensity of ASCP and reduction of ThT fluorescence observed in the binding-competition assay suggests that the binding sites of ASCP to α-synuclein fibrils are the same or overlap with the binding sites of ThT. Molecular structures obtained through NMR, crystallography and super-resolution imaging have provided insights into the apparent binding mechanism of ThT to amyloid structures ^47–49^. Since amyloid fibrils are diverse in their underlying amino-acid sequence, it is generally accepted that ThT recognises structural features common to all amyloids, in particular, the continuous cross-β-sheet structures and side-chain arrangements known as cross-strand ladders ^9^. Amyloidogenic dyes, including ThT and Congo Red, have been shown to bind to the β-sheet surface, orientated within the channel-like motifs formed by the cross-strand ladders ^50^. As ASCP is structurally similar and competitively displaces ThT molecules bound to α-synuclein fibrils it is suggested that ASCP also binds within the cross-strand ladders of the β-sheet structures of α-synuclein fibrils; however, subsequent NMR and crystallography experiments would be required to confirm this hypothesis.

The ability to perform *in situ* assays to monitor the fibrillar aggregation of proteins is particularly advantageous, as key kinetic and mechanistic processes can be monitored in real-time ^21,28,29^. Thus, we sought to monitor the real-time aggregation of RCM κ-casein, a well characterised amyloid-forming protein ^51,52^, as well as the aggregation of the Aβ_1-42_ and *α*-synuclein proteins, which are associated with the pathogenesis of Alzheimer’s and Parkinson’s disease, respectively ^53–55^. Interestingly, our results suggest that ASCP is more sensitive to oligomeric species formed early during aggregation as there tended to be an earlier and more pronounced increase in ASCP fluorescence observed when monitoring the aggregation of RCM κ-casein and Aβ_1-42_ compared to ThT. Similar results have been observed previously for another aggregation-induced emission fluorophore, TPE-TPP, which was shown to be significantly more sensitive to oligomeric species formed early during the fibrillar aggregation of RCM κ-casein ^22^. Similarly, ASCP appears to report on the formation of species, such as small oligomers and fibril nuclei, present early during the aggregation of Aβ_1-42_, including the period typically described as a lag phase when ThT is used in these assays. In contrast, an increase in ThT fluorescence was not observed until later times, corresponding to the elongation phase of the reaction.

Indeed, one of the main limitations of ThT is its relative inability to detect oligomeric species formed early during aggregation ^7^, which is of particular importance with recent evidence suggesting that these oligomers are the primary cytotoxic species responsible for the pathogenesis of numerous aggregation-associated diseases ^56^. The work presented here demonstrates that ASCP has a superior ability to detect these soluble, pre-fibrillar aggregates compared to ThT and therefore ASCP may be more suited for the real-time monitoring of protein aggregation. Importantly, since lower concentrations of ASCP did not significantly change the fluorescence profile associated with Aβ_1-42_ fibril formation, it can be inferred that ASCP does not interfere with the kinetics of aggregation and that the dye is likely reporting on nucleation events that are not able to be visualised using ThT.

Interestingly, it was found that the ability of ASCP to monitor the aggregation of proteins was not strictly limited to the detection of amyloid fibrils since significant increases in fluorescence were also observed when it was present during the amorphous aggregation of *α*-lactalbumin and insulin. The finding that ThT was also responsive to amorphous aggregation was surprising since an increase in ThT fluorescence has long been associated with the specific detection of amyloid aggregates. The increase in fluorescence of both of these dyes in the presence of amorphous aggregates likely represents their generic mechanism of fluorescence, whereby restriction of intramolecular rotations is the only pre-requisite for increased fluorescence emission ^7^. Both dyes, for instance, have been shown to be fluorescent in the presence of other biological macromolecular structures that do not contain the β-sheet structure characteristic of amyloid fibrils, such as those formed by DNA and RNA ^24,57–59^. Thus, it is likely that both ThT and ASCP can report on amorphous aggregation by binding to hydrophobic grooves or cavities large enough to accommodate the dyes and restrict their intramolecular rotations. It is possible that the relatively small size of these dyes facilitates their binding to amorphous aggregates, since larger AIE fluorogens such as TPE-TPP, which is twice as large as ThT, do not possess the ability to monitor the amorphous aggregation of proteins ^22^. Interestingly, ASCP appears to be able to monitor the early stages of the amorphous aggregation of *α*-lactalbumin in a manner similar to that observed for the fibrillar aggregation of RCM κ-casein and Aβ_1-42_, which again suggests that ASCP is able to report on the formation of small, oligomeric species not large enough to be detected by light scatter or ThT. Together, the results of the competitive binding and aggregation assays suggest that ASCP not only shares a binding site with ThT but may also possess an additional binding mode/s compared to ThT that enables ASCP to detect early oligomeric species formed early during both amorphous and fibrillar aggregation.

Data from this study and others have demonstrated that high concentrations of ThT (e.g. 50 µM) do not significantly affect the kinetics of fibril formation ^60^. Consequently, many real-time and *ex situ* fluorometric assays have been developed that utilise ThT to monitor fibril formation ^14–16,18,61^. However, caution must still be employed to ensure that the addition of an extrinsic probe to the fibrillation reaction does not interfere with the aggregation kinetics. In this work we demonstrate that ASCP can be used at concentrations of up to 5 µM to monitor the elongation of *α*-synuclein fibrils *in-situ* via increases in fluorescence intensity over time. However, ASCP was found to reduce the elongation rate of *α*-synuclein fibrils at concentrations greater than 5 µM, and the total fluorescence and amyloid content at concentrations greater than 10 µM.

This inhibitory effect of ASCP mirrors the *in situ* inhibition of amyloid fibril elongation observed with other dyes and amyloidogenic proteins, including a tetraphenylethene ^62^ and Congo Red ^63^. Certain polyphenols and other stilbene derivatives similar to ASCP are established inhibitors of amyloid formation, including curcumin ^64,65^ and resveratrol ^66,67^. The suggested mechanism of inhibition is the binding of the fluorogenic dye to an exposed hydrophobic region in partially unfolded protein monomers ^62,68^. This interaction affects the formation of the cross-β-sheet motifs and prevents the ‘docking’ or incorporation of the monomers onto the fibril ^62,69^. Therefore, the additional aromatic structure present in ASCP, compared to ThT, may enable ASCP to form more favourable hydrophobic interactions that prevent monomer ‘docking’, contributing to the inhibitory effects of ASCP on *α*-synuclein fibril elongation at concentrations above 10 μM. Regardless, the results of this work have demonstrated that it is possible to use ASCP to monitor the *in situ* elongation of fibril elongation, albeit under optimised conditions and lower concentrations of dye relative to ThT. Moreover, we show ASCP is ideal for *ex situ* investigations of fibril formation. Furthermore, ASCP was not found to interfere with the aggregation of other proteins used in this study and thus the observed amyloid inhibition effect observed for *α*-synuclein fibril elongation at high concentrations of ASCP may be a protein-specific phenomenon.

Aggregation assays that employ the use of aggregation-sensitive fluorophores, such as ThT, have been extensively employed to study the kinetics of fibril growth (e.g. lag and elongation rates); however, the underlying mechanisms of amyloid formation are complex and extremely heterogeneous. Thus, single-molecule approaches that enable the characterisation of individual amyloid fibrils are becoming increasingly common ^14–18,61,70^. The use of AIE fluorophores has a substantial advantage in single-molecule investigations of amyloid formation since photobleaching is avoided and the protein of interest does not need to be covalently labelled with a fluorescent probe, which can change the kinetics and overall morphology of the elongating fibril ^17,61,70–74^. With this in mind, we sought to investigate if ASCP could serve as a suitable alternative to ThT in single-molecule TIRF microscopy experiments. The results of this study found that *α*-synuclein fibrils stained with relatively low concentrations of ASCP could easily be visualised using TIRF microscopy and the length of these fibrils was observed to increase throughout incubation. Furthermore, as a proof of principle, we demonstrate that the molecular chaperone *α*B-c can bind to pre-formed *α*-synuclein fibrils and that both species can be visualised simultaneously when illuminating the sample with a single 488 nm excitation laser source.

There are several advantages in using ASCP in microscopy experiments compared to ThT. First, the significant Stokes shift of ASCP allows a single illumination source to simultaneously excite ASCP and another fluorophore (e.g. Alexa Fluor 488) without the need to worry about fluorescence spectral overlap, which facilitates the study of more complex biological systems without additional technical complexity. The spectral separation required for single-laser multi-fluorophore imaging simply cannot be achieved with probes typically used for biological applications (i.e. ThT, GFP and Alexa Fluor dyes) due to their small Stokes shift. Second, dyes that exhibit large Stokes shifts (> 100 nm) also reduce light scattering and interference between the excitation source and fluorescence emission detection ^75,76^, resulting in significantly improved signal to background ratios and enhanced localisation of individual fluorophores. Furthermore, the use of dyes such as ThT and TPE-TPP, which emit in the blue region of the spectrum, is severely restricted in experiments involving cells or tissues due to interference of the dye signal as a result of autofluorescence. The significant red-shifted emission of ASCP overcomes such limitations and with further development could be used for the study of protein aggregates in cells and tissues. Finally, the excitation properties of ThT (412 nm) are sometimes inaccessible to conventional filter setups of laboratory instruments, including flow cytometers and fluorescence microscopes, which are typically equipped with 488 nm lasers as standard. Therefore, ASCP has the advantage of being more compatible with standard optical configurations (488 nm/647 nm), that are most common in fluorescence-based instruments.

## Conclusions

The use of extrinsic dyes, such as ThT, to detect and monitor the growth of amyloid fibrils *in vitro* has become commonplace and has greatly contributed to our understanding of the pathology and progression of aggregation-associated diseases. ThT is the most utilised and characterised dye for the detection of amyloid; however, there is a growing need to develop alternative dyes that possess an improved ability to detect soluble or pre-amyloid species formed early during aggregation and have unique and enhanced spectral qualities compared to ThT. To this end, we investigated the photophysical and fibril-binding properties of the *α*-cyanostilbene derivative ASCP via absorbance and fluorescence spectroscopy as a potential alternative to ThT. Similar to ThT, ASCP became fluorogenic in the presence of amyloid fibrils, exhibited a higher binding affinity to *α*-synuclein fibrils compared to ThT and also competitively displaced bound ThT from *α*-synuclein fibrils that suggests a shared binding site. Furthermore, ASCP can be used to monitor amorphous and fibrillar aggregation when used *in situ* and has a superior ability to detect soluble oligomers or pre-fibrillar species compared to ThT. Interestingly, ASCP also exhibits a pronounced Stokes shift of 145 nm that can be utilised in TIRF microscopy experiments to simultaneously visualise fluorescently labelled *α*B-c chaperone and ASCP-stained *α*-synuclein fibrils using only a single laser. Consequently, we propose that the red-shifted emission, enhanced Stokes shift and fibril-binding properties makes ASCP a superior alternative to ThT for the study of amyloid fibrils in TIRF microscopy experiments.

## Author contributions

HE, AvO and YH initiated the work. YH supplied the ASCP dye. NRM, CLJ, BPP, HE and AvO formulated the experimental approach. CLJ, KMW, NRM and BPP developed the conditions for the TIRF experiments. NRM, KMW performed all other experiments. NRM, KMW, and CLJ collated the data and constructed the figures. NRM, KMW and HE wrote the manuscript. All authors edited the manuscript and approved the submission of the final version.

## Acknowledgements

This research performed by NRM and CLJ has been conducted with the support of the Australian Government Research Training Program Scholarship. AvO is supported by an ARC Laureate Fellowship (FL140100027). We thank Tze Cin Owyong and Anuradha for synthesis of the dye. Finally, we thank staff in Molecular Horizons and the Illawarra Heath and Medical Research Institute for technical and administrative support.

