## Supplementary Information for "An *α*-cyanostilbene derivative for the enhanced detection and imaging of amyloid fibril aggregates"

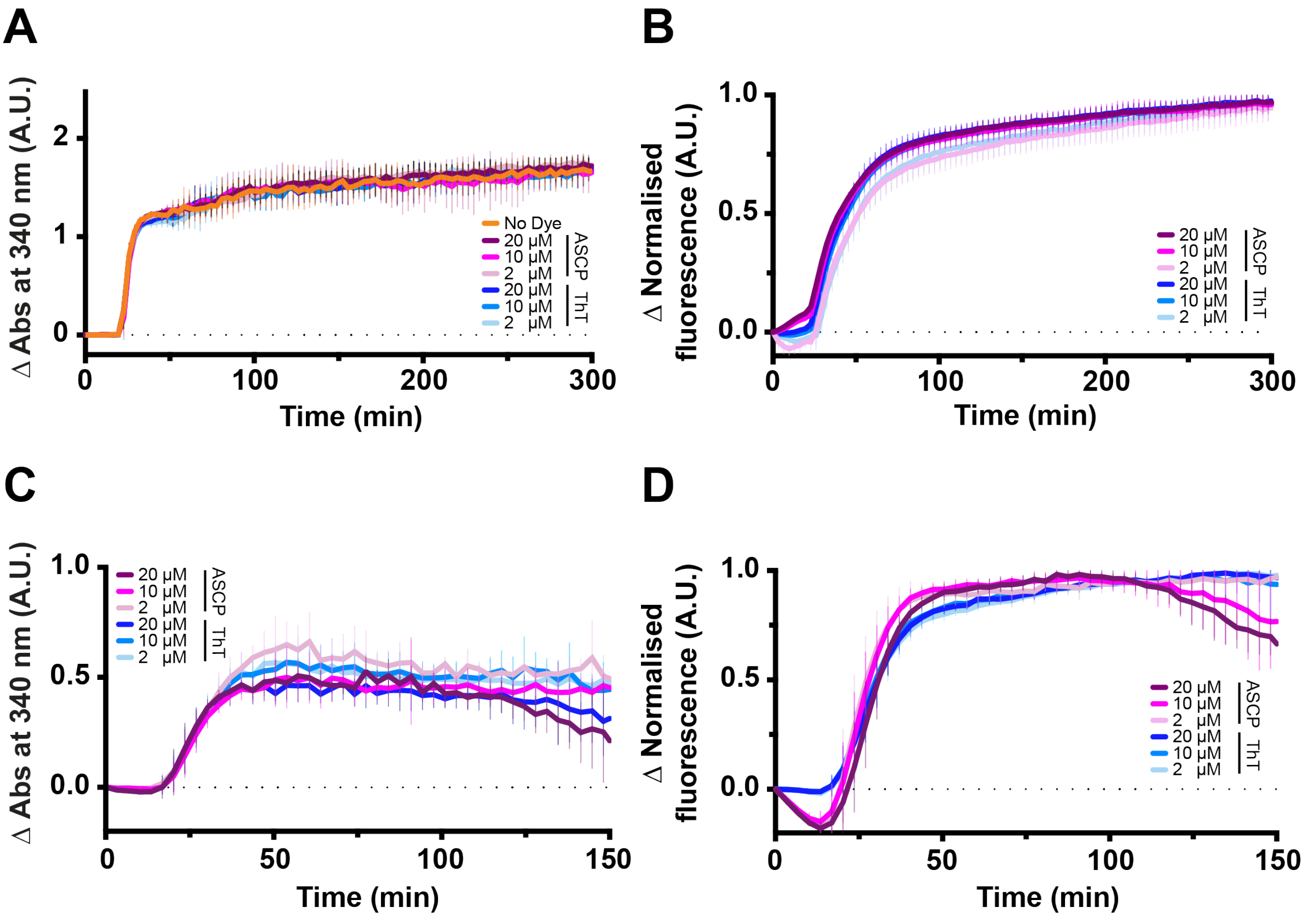


**Supplementary Figure 1: ASCP does not affect the kinetics of amorphous aggregation *in situ*.** Various concentrations of ThT or ASCP (0 - 20 µM) were added to $\alpha$-lactalbumin (100 µM, **A and B**) or insulin (100 µM, **C and D**) and the aggregation of protein was monitored via absorbance at 340 nm (A and C) and fluorescence (B and D). The fluorescence was measured using excitation/emission wavelengths of 440/490 nm and 485/590 nm for ThT and ASCP, respectively. $\alpha$-lactalbumin and insulin aggregation assays were run in triplicate and presented as the mean ± standard deviation from at least 3 independent experiments.

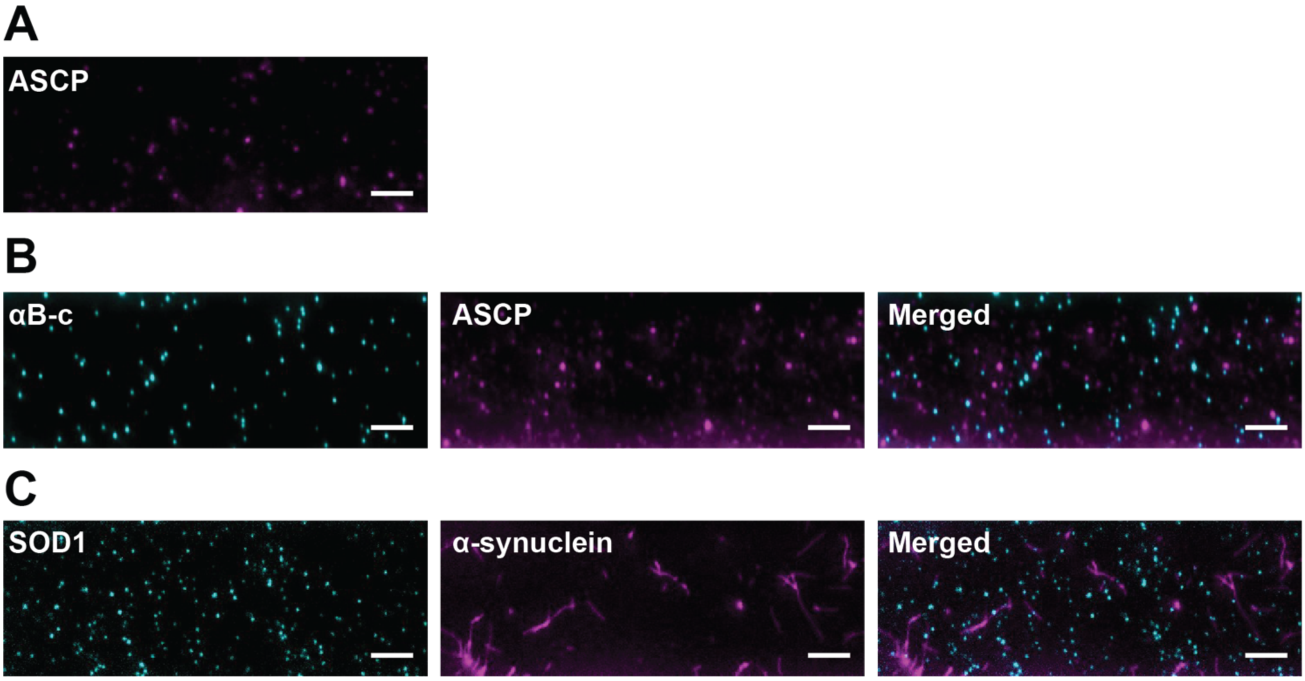


**Supplementary Figure 2: Control experiments for the use of ASCP in single-laser two-colour TIRF microscopy experiments.** (**A**) A representative TIRF image of $\alpha$-synuclein monomer (50 µM) that was diluted 4,000-fold in in 20 mM phosphate-trolox buffer (pH 7.4) containing ASCP (5 µM). (**B**) AF488-$\alpha$B-c (1 µM) was incubated in 20 mM phosphate-trolox (pH 7.4) for 30 min at room temperature in the absence of $\alpha$-synuclein and diluted 1,000-fold in buffer containing ASCP (5 µM). (**C**) Fibrillar $\alpha$-synuclein (50 µM) was incubated in 20 mM phosphate-trolox buffer (pH 7.4) in the presence of 1 µM AF488-SOD1 for 30 min at room temperature. Samples were diluted 1,000-fold into buffer containing ASCP (5 µM). Samples from B and C were imaged using TIRF microscopy and the fluorescence emission of AF488-labelled species and ASCP were collected in separate channels. Representative TIRF images for each experiment are shown. The scale bars represent 5 µm.
